# The susceptibility analysis of *Echinococcus multilocularis* protoscoleces tubulin to mebendazole and RNA interference

**DOI:** 10.1101/2020.09.24.312736

**Authors:** Qiqi Shi, Lele Huo, Bin Jiang, Haijun Gao, Jinxin Zheng, Yi Tao, Haobing Zhang

**Affiliations:** National Institute of Parasitic Diseases, Chinese Center for Disease Control and Prevention, Chinese Center for Tropical Diseases Research, WHO Collaborating Centre for Tropical Diseases, National Center for International Research on Tropical Diseases, Ministry of Science and Technology, Key Laboratory of Parasite and Vector Biology, Ministry of Health, Shanghai, China

**Keywords:** mebendazole, RNAi, tubulin, flame cell, *Echinococcus multilocularis*

## Abstract

Alveolar echinococcosis, caused by the larval (metacestode) stage of the tapeworm *Echinococcus multilocularis*, is a lethal parasitosis of the liver prevalent in the Northern Hemisphere. For chemotherapy the benzimidazole derivatives mebendazole and albendazole were introduced, which were found to disrupt the microtubules by inhibition of the polymerization of tubulin into microtubules, and β-tubulin was determined to be the drug target molecule. In the present study, we evaluated the chemosensitivity of *E. multilocularis* protoscoleces tubulin to mebendazole and RNA interference in vitro, and to explore whether the molecular level and ultrastructure of *E. multilocularis* protoscoleces microtubules change post-mebendazole and RNA interference. We identified that mebendazole is parasitostatic to *E. multilocularis* protoscoleces through suppression the tubulin expression and change the flame cell morphology in molecular level, besides RNA interference indicated that *β 2* tubulin is probably one of the vital tubulin gene to form the flame cell and the protonephridial system tubules (collective tubes) of *E. multilocularis* protoscoleces. Molecular level and ultrastructure detection were performed by reverse transcription-PCR, western blotting and transmission electron microscope. The RNA interference would be probably as a parasitocidal method to disrupt the survival of PSCs, extend that the relevant tubulin maybe as potential target for drug development against AE.

## INTRODUCTION

The larval stage of the fox tape worm, *Echinococcus multilocularis*, is the causative agent of alveolar echinococcosis (AE), which is considered to be the most lethal human helminthiasis (1). In 2014, the Food and Agriculture Organization of the UN and WHO, based on a multicriteria ranking system, highlighted AE and CE (cystic echinococcosis) as the second and third most important food-borne parasitic diseases at the global level, respectively (2). The prevalence of CE and AE extends beyond tropical and subtropical regions to include worldwide pastoral and rural communities of medium-high income countries, including China, the Russian Federation, Europe and North America. In Europe, where they should be managed as orphan diseases (3,4). CE and AE combined infect more than 1 million people worldwide at any given time, with an estimate of 200 000 new cases per year (5). In central Asia, 270 million people are at risk of CE or AE, which, after the collapse of the Soviet Union and the consequent profound socioeconomic changes, are considered to be re-emerging in some countries, such as Kyrgyzstan, Tajikistan, Turkmenistan, and Uzbekistan (6). During the period between 1995 and 2011, human AE in Kyrgyzstan increased at least 20-fold, from 0–3 cases to more than 60 cases per year (7). The disease burden of selected zoonotic infections in Kyrgyzstan (AE, brucellosis, campylobacteriosis, CE, congenital toxoplasmosis, rabies, and non-typhoidal salmonellosis) is substantial and similar to that of HIV (35 209 *vs* 38 870 disability-adjusted life-years [DALYs] in 2013) (8). Among these zoonotic agents, the second highest burden was caused by *EM* (11 915 DALYs [95% uncertainty interval 4705–27 114] per year) (8).

Human AE is characterized by an asymptomatic incubation period of 5–15 years (progressing infiltrative proliferation of the metacestode into adjacent organs and tissues) and the slow development of a primary tumour-like lesion which is usually located in the liver, with a high fatality rate if untreated, which the mortality rate in humans is >90% (9). The only curative treatment for AE is complete surgical resection of the parasite tissue. Such invasive surgery is performed in about 30% of all AE patients, therefore most receive only continuous medication with the benzimidazole-derivatives albendazole (ABZ) or mebendazole (MBZ) (10). Benzimidazoles have drastically improved the life expectancy and quality of life of patients. Whereas the 10-years survival rate of untreated AE patients was 0–25% in the pre-benzimidazole era, benzimidazole-treated patients to date have a 10-years survival rate of 91–97% in countries with well-developed healthcare (11,12).

Several studies performed with AE patients have shown the benefit of chemotherapy with MBZ, increasing the survival time of the patients, providing an improved clinical condition, and decreasing the size of the parasitic mass (13–16). However, limited data on experience with therapy with ABZ are available (17). In other aspect, in immunocompromised patients, reducing immunosuppressive treatment must be considered when this is possible. In the French cohort, ABZ efficacy was shown to be fast and excellent but with more adverse effects than in non-immunocompromised patients treated in the same centers (18).

Benzimidazole (BZ) compounds were found to disrupt the microtubules by inhibition of the polymerization of tubulin into microtubules, and β-tubulin was determined to be the drug target molecule (19, 20). The selective activity of BZ mediated mainly against the parasite, is due to higher affinity of the BZ to the parasite tubulin compared to that of the host (21). Microtubules are cylindrical structures in eukaryotic cells that mediate many processes including mitosis, ciliary and flagellar motility, and intracellular transport of vesicles and organelles. In addition, they are the most common component of the cytoskeleton, governing cell morphology (22). In addition to β-tubulin, α-tubulin is a major building block of microtubules. These two components are similar in mass and form a heterodimer (23). In most eukaryotes, α- and β-tubulin undergo various post-translational modifications; therefore, they exist as families of related isoforms (24). However, because until now no susceptibility testing of the *E. multilocularis* metacestodes tubulin to BZ derivatives in vitro, it has remained uncertain which tubulin gene isoform was susceptible to the BZ derivatives, and which tubulin gene isoform was pivotal and significant to metacestodes development. To address these problems, we used a recently established in vitro system to analyze the action mode of MBZ and RNAi against *E. multilocularis* tubulin from molecular level.

## MATERIALS AND METHODS

### Biochemicals

Mebendazole was purchased from Sigma-Aldrich (Buchs, Switzerland).

### *In vitro* culture of *E. multilocularis* metacestodes and viability assessment

*E. multilocularis* -infected animals [BALB/c mice (SLAC Laboratory Animal Center)] were maintained by serial intraperitoneally (i.p.) transplantation passages through for 6 months, the details can be found in our published paper (25). Eighty infected BALB/c mice were prepared for *in vivo* treatment experiments. Parasite materials were isolated from the peritoneal cavity of the infected animals and homogenized with Hanks’ balanced salt solution (HBSS). Then, after passing the parasite homogenate through a 60-mesh sieve, the *E. multilocularis* PSCs were collected and rinsed 5~8 times. The viability of the PSCs was determined by the methylene blue exclusion method (26), and PSCs with > 95% viability were used in the following experiments. The PSCs were cultured in RPMI 1,640 medium supplemented with 10% fetal bovine serum and antibiotics (100 U/ml penicillin G and 100μg/ml streptomycin) at 37° C in 5% CO2 (27), before distribution in 24-well plates (1 ml vesicle suspension per well, about 2000 metacestodes per well).

### *In vitro* drug treatment of *E. multilocularis* metacestodes

The metacestodes after 1 day of culture in vitro, MBZ were pre-diluted in DMSO at the working concentration of 10mM, 5mM and 2.5mM, then respectively added to the wells (5ul per well) to fixed final concentrations at 50uM, 25uM and 12.5uM, every concentration in triplicate. Corresponding amounts of DMSO (5ul) were used as the negative control in triplicate. The parasite- and drug-containing plates were incubated at 37 °C and 5% CO2, under humid atmosphere. To assess drug-induced metacestode tubulin damage by qRT-PCR, the mixture of each well was collected after 1, 3, 5 and 7 days of incubation, then stood in TRIzol reagent (Invitrogen, CA, USA) for 15min, following stored at −80 °C until further total RNA extracted were performed.

### *In vitro* drug testing of *E. multilocularis* metacestodes in mRNA level

The Efficacy analysis of MBZ in mRNA level were applied with qRT-PCR. MBZ-treated metacestodes were collected at 1 day, 3day, 5day and 7 day. Total RNA was extracted from PSCs using TRIzol reagent (Invitrogen, CA, USA), according to the manufacturer’s instructions. The quality and concentration of RNA was confirmed and 2ug of the RNA sample was treated with DNase I (Thermo, Waltham, MA, USA) for 30 min at 37 °C to remove genomic DNA contamination (28). The cDNA was synthesized from the treated RNA using Primer Script RT kit (Takara Bio, Dalian, China), according to the manufacturer’s instructions. qRT-PCR was performed for gene expression using 2uL of 1:5 diluted cDNA with SYBR Green Realtime PCR Master Mix and Permix Ex Taq (Takara Bio), according to the manufacturer’s instructions. The sense and antisense primers for *E. multilocularis* tubulin *α9, β2, β4, β6* and *EF-1α* were synthesized by Sangon Biotech (Shanghai, China). The sequence information was as follow (Table 1).

**Table 1:**
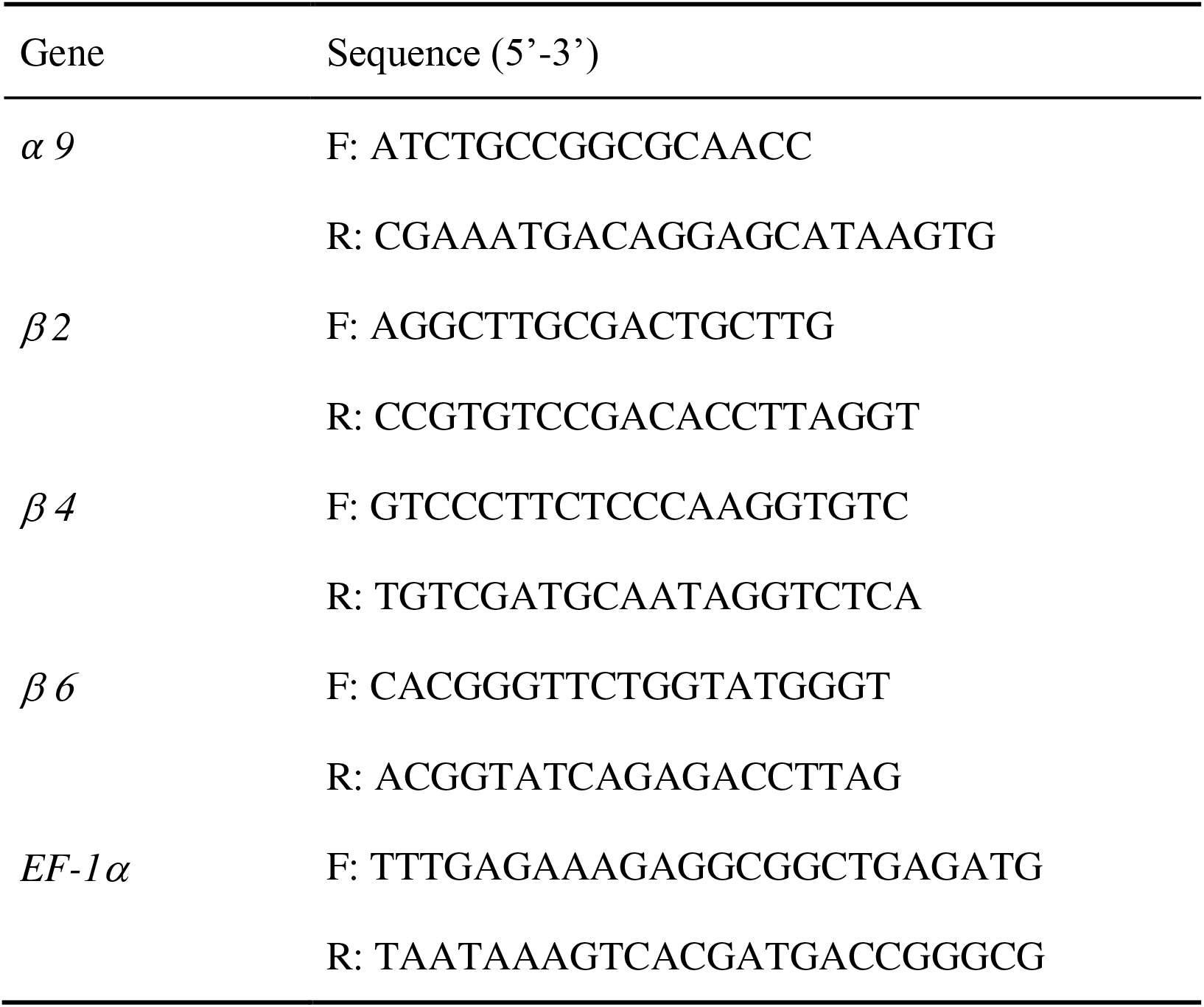
The primer sequence information of *α9, β2, β4, β6 and EF-1α*. for qPCR.

qRT-PCR was operated on ABI Prism 7500 Sequence Detection System (BioRad, Life Science Research, Hercules, CA, USA). All samples were performed in triplicate using the following cycle parameters: one cycle at 95°C for 30 s, 39 cycles at 95°C for 5 s, at 60°C for 10s, at 72°C for 30s. A melting point curve was analysed after the PCR by increasing the temperature from 65 to 95 °C (0.5 °C increments) to validate the PCR amplification specificity. Furthermore, the PCR amplification efficiency of the target gene tubulin *α9, β 2, β 4, β 6* (all the E were greater than 104.7%, R were greater than 0.994) and reference gene *EF-1α* (E = 108.3%, R^2^ = 0.993) was established by standard curves. The 2^−ΔΔCT^ method was used to calculate relative concentration of each target by standardizing to internal *EF-1α* level. Each relative value was normalised to the DMSO treated PSC sample.

### *In vitro* drug testing of *E. multilocularis* metacestodes in protein level

The efficacy analysis of MBZ on protein levels were completed by western blotting. The treated PSCs were collected on day 1, 3, 5 and 7 after administration. Proteins were isolated from PSC samples used T-PER™ Tissue Extraction Reagent (Invitrogen), according to the manufacturer’s instructions. Protein concentration was measured using the BCA protein assay. We use 4-12% (w/v) NuPAGE Bis-Tris protein gels (Invitrogen precast gel, Invitrogen) including 15 wells with the Mini Blot module in a Mini Gel Tank (Invitrogen) at 150 V for 45 min at room temperature to separate proteins. We routinely run whole-well gels, loading 10μg protein per gel. Then, the protein is transferred onto polyvinylidene fluoride (PVDF) membranes (Millipore Corp., MA, USA) at 20 V, 60min at room temperature. The membrane was washed with Tris buffered saline with Tween 20 (TBST) [G-Biosciences] and blocked with LI-COR/PBS buffer (LI-COR Biosciences, Lincoln, NE, USA) for 1h at room temperature, then incubated with the monoclonal mouse anti-α-tubulin IgG, rabbit anti-β-tubulin IgG (Cell Signaling Technology, Danvers, MA, USA) in a 1:1000 dilution with LI-COR/PBS buffer denatured in 4°C overnight. The internal reference protein was GAPDH protein, and antibody was rabbit anti-GAPDH IgG (Invitrogen, Carlsbad, CA, USA) in a 1:2000 dilution, all is HRP-conjugated. Bands were visualized using enhanced chemiluminescence (ECL) with ChemiDoc XRS+ (bio-rad, America) system.

The qRT-PCR and western blotting analyses were performed in triplicate for each group. Each treatment was carried out in triplicate and the whole experiments were repeated twice using samples of PSCs collected at different times.

### Ultrastructural investigations of MBZ-treated metacestodes

To visualize the structural alterations in metacestodes imposed by MBZ treatment, parasites were processed for transmission electron microscopy (TEM) at day 7 after the initiation of treatment with 50uM, 25uM, 12.5uM MBZ. Briefly, the steps of specimen preparation for TEM were carried out as described by Küster T, et al (29) and finally, 70-nm thick sections were imaged on a transmission electron microscope at 80 kV (JEOL JEM-1400) using a TempCam F416 camera (Tietz Video and Image Processing Systems GmbH, Gauting).

### *In vitro* RNA interference treatment of *E. multilocularis* metacestodes

To optimize the transfection effect, two methods, soaking and electroporation, were used to deliver a Cy3-labelled (red fluorescence) siRNA into parasites. As expected, fluorescence was not detectable in the untreated PSCs. The soaking method resulted in a relative high level of death rate as only PSCs appeared black and shrink in the tegument of PSCs. In contrast, electroporation resulted in high efficacy with the average transfection efficiency being 97%, nearly all PSCs were visualized the resulting emitted fluorescent light and have no death occur (Fig. 1A). According to the reference (30), We pre-tested the *elp* and *14-3-3* siRNA, which were Fam-labelled (green fluorescence), to further verify the process of electroporation and determine the positive control. The electroporation of *14-3-3*-siRNA and *elp*-siRNA were also feasible (Fig. 1B). The experiment was repeated three using PSC samples collected at different times.

**Fig 1.**
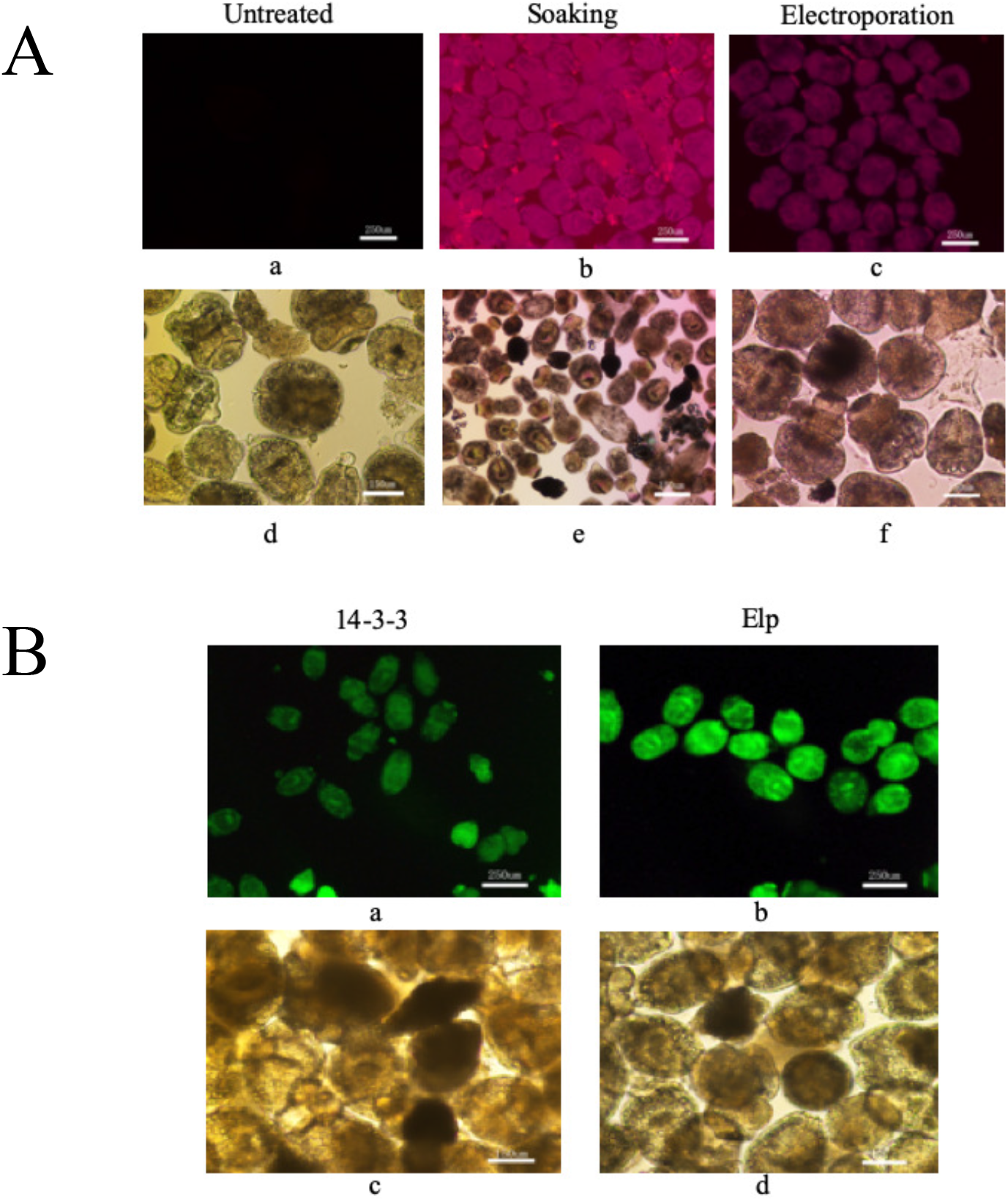
A showed transfection of siRNA into PSCs. Localisation of Cy3-labelled (red fluorescence) siRNA in PSCs following soaking or electroporation at 2 h was observed by confocal microscopy. B showed transfection of *14-3-3*-siRNA and *elp*-siRNA into PSCs through electroporation method. Localisation of Fam-labelled (green fluorescence) siRNA in PSCs following electroporation at 2 h was observed by confocal microscopy. The scale bar represents 250 um at A (a, b, c) and B (a, b); while the scale bar represents 150 um at A (d, e, f) and B (c, d).

The pre-test result (qRT-PCR) suggested that the *14-3-3* gene was knocked down (the *14-3-3* zeta mRNA reduced 12.82 ratio in day 3, lower than the levels of the untreated control, while *elp* siRNA-treated sample was reduced 3.78 ratio than that of untreated control), so we applied *14-3-3* siRNA as the positive control. We designed and synthesized 6 fragments of siRNA targeting tubulin *α 9, β 2, β 4, β 6* genes from Sigma-Aldrich at a concentration of 100μM. Nine sample groups of PSCs for electroporation were set: untreated controls, positive controls (*14-3-3* siRNA-treated), *α 9-*siRNA-treated samples, *β 2-1*-siRNA-treated samples, *β 2-2*-siRNA-treated samples, *β 4*-siRNA-treated samples, *β 6-1*-siRNA-treated samples, *β 6-2*-siRNA-treated samples, *β 6* t-siRNA-treated samples (*β 6-1*-siRNA and *β 6-2*-siRNA treated the sample jointly). At 30 min and 2 h after treatment, the parasites were washed with PBS and viewed under a fluorescent microscope (FV500, Olympus) to evaluate the efficacy. Then the successful treated PSCs were incubated for a further two weeks in RPMI 1640 (Gibico) supplemented with 10% (v/v) heat-inactivated FBS (Gibico), 100 U/mL penicillin, and 100 ug/mL streptomycin (Hyclone) at 37 °C in an atmosphere of 5% CO2 in dark, for further verified the RNAi effects from mRNA and protein levels. During this period, approximately half of the medium was changed every 3 days. Each treatment was carried out in triplicate and experiments were repeated twice using samples of PSCs collected at different times. The sequence of the 6 fragments is described in Table 2.

**Table 2:**
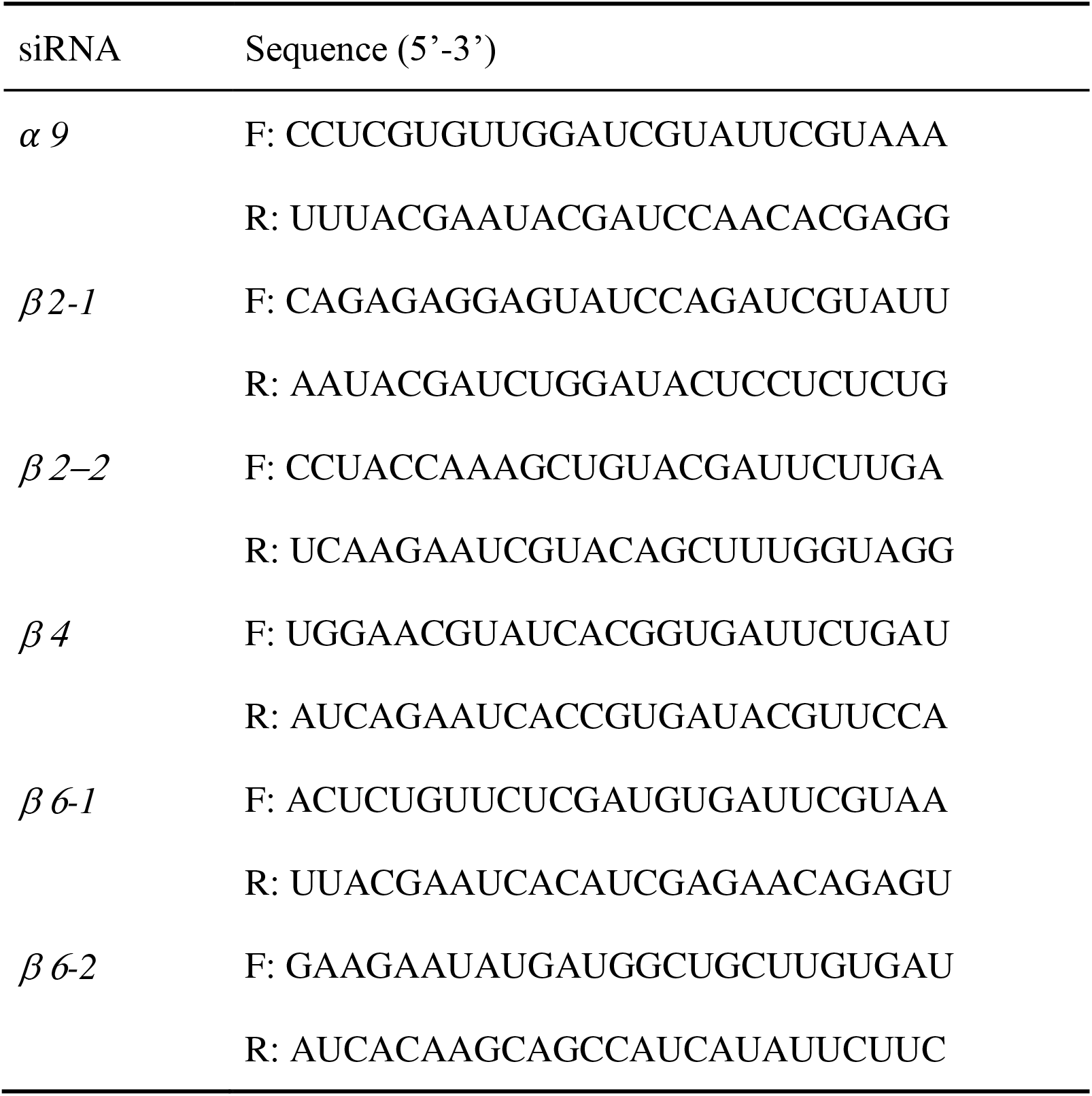
The sequence information of *α 9, β 2, β4, β 6* siRNA fragments for RNAi.

### *In vitro* efficacy determination of *E. multilocularis* metacestodes post-RNAi in mRNA level

The efficacy of RNAi on mRNA levels were evaluated using qRT-PCR. For the analyses, total RNA was first extracted from all groups at day 3 after electroporation using TRIzol reagent (Invitrogen), as previously described, and the primer sets as follow: *14-3-3 zeta* forward 5’-AAC TTG CTA TCC GTT GC-3’, reverse 5’-CAC CTT CTT AAG GTA AAT GTC-3’; *elp* forward 5’-GTG AAG TCT GGT ACT TCG-3’, reverse 5’-ATC CAG TCT TAG AAA GGT TG-3’(31). *EF-1 a* was as above. The residual primer information was same with above.

### *In vitro* efficacy determination of *E. multilocularis* metacestodes post-RNAi in protein level

The efficacy of RNAi on protein levels were evaluated using western blotting. The treated PSCs were collected on day 6, 10, and 15 after electroporation and homogenized using a hand homogenizer (23M-R25, Nippon Genetics). The assay was accomplished as previously described. The primary antibodies were rabbit anti-14-3-3 (31), rabbit anti-antigen II/3 (32) antibodies, and the goat anti-rabbit antibody (anti-rabbit IgG AP conjugate, Promega) as secondary antibody (30). The internal reference protein was GAPDH protein, and antibody was rabbit anti-GAPDH IgG HRP-conjugated (Invitrogen, Carlsbad, CA, USA) in a 1:2000 dilution.

The qRT-PCR and western blotting analyses were performed in triplicate for each group. Each treatment was carried out in triplicate and the whole experiments (including the following effect verify) were repeated twice using samples of PSCs collected at different times.

### Ultrastructural investigations of RNAi-treated metacestodes

To visualize the structural alterations in metacestodes imposed by RNAi treatment, parasites were processed for TEM at day 6, 10, 15 after the electroporation treatment with siRNA. We tested untreated controls samples, *β 2-1* -siRNA-treated samples and *β 2-2*-siRNA-treated samples. The test was carried out as previously described.

### Statistical Analysis

Statistical analysis was performed using GraphPad Prism 8.3.1 software (San Diego, CA, USA). Data were expressed as the mean ± standard error of the mean and analysed statistically using one-way ANOVA with Tukey’s multiple comparison test at each time point was performed to determine differences between four groups. *p* < 0.05 was regarded as statistically significant.

## RESULTS

### *In vitro* efficacy of MBZ against *E. multilocularis* metacestodes tubulin

As illustrated in Fig. 2, the fold change (FC) value of *α9, β 2, β 6* tubulin genes on day 1 were less than 1, subsequently were greater than 1, revealed that these genes appeared higher expression when MBZ stimulate originally, however, with time extend these genes had lower expression. In day 3 group, MBZ significantly (*P* ≤ 0.05) inhibited the intracellular replication of *α 9, β 2, β 4, β 6* tubulin genes at the concentration range of 12.5–50uM. In day 7 group, the FC value of *α 9, β 2, β 4, β 6* tubulin genes under 25uM was larger than 2, suggested that these four tubulin genes were down expressed approximately in a time course manner under 25uM concentration. However, in day 1, 5 and 7 groups, there were no significant differences between different concentration, as well different genes.

**Fig 2.**
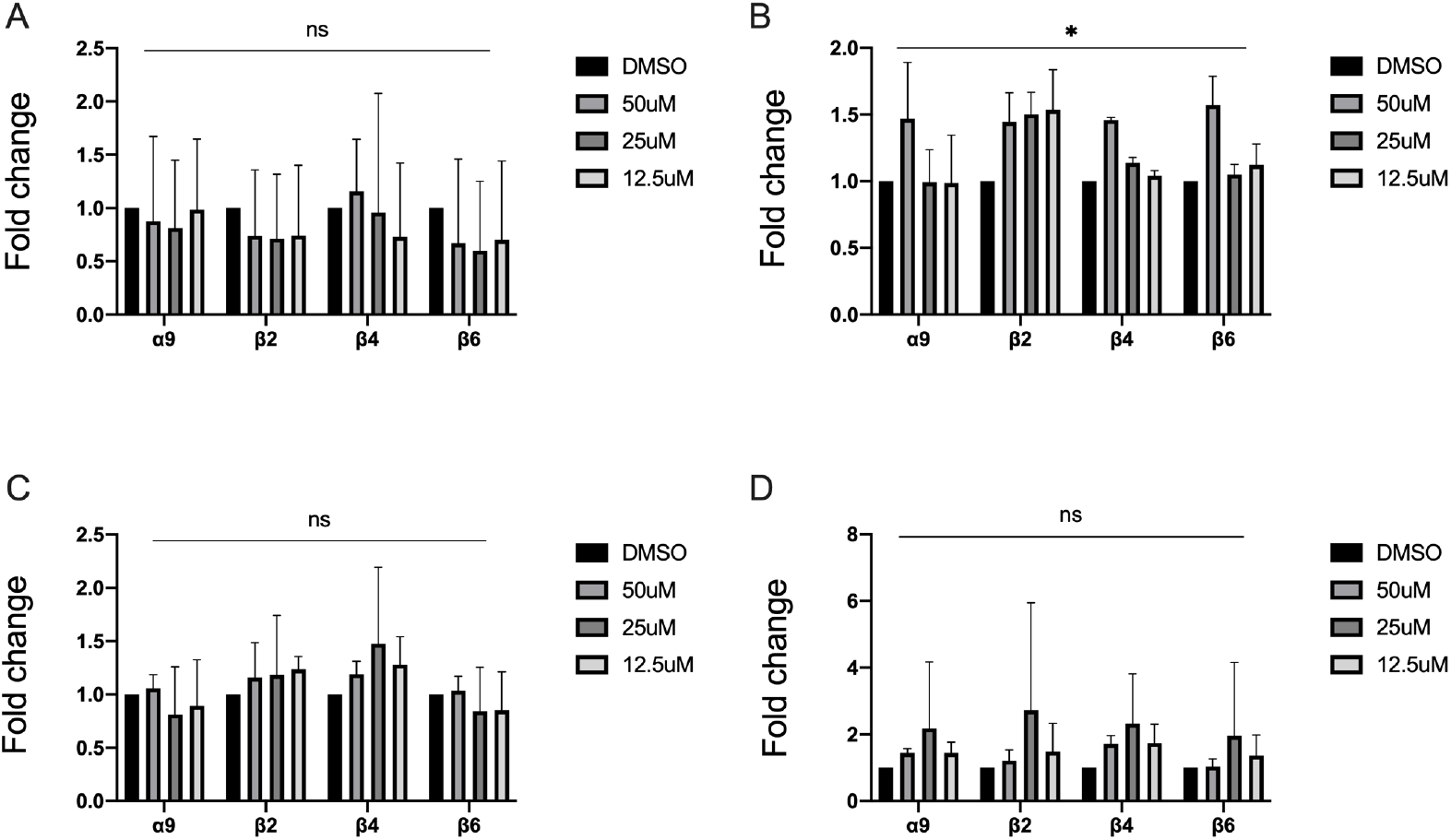
Quantitative PCR analysis of tubulin mRNA in different concentration-treated or DMSO-treated PSCs by day1, day3, day5 and day7 (A, B, C, D represents day1, 3, 5, 7 respectively). Bars represent the standard deviation. *p < 0.05. ns: no significance.

As shown in Fig. 3, the FC value of *α 9, β 2, β 4* tubulin genes were gradually increased with time and concentration increment. This reveled that these tubulin genes changes over administrate time and concentration, furthermore, 25uM concentration of MBZ after 7 day maybe showed the optimized tubulin-targeting efficiency in vitro.

**Fig 3.**
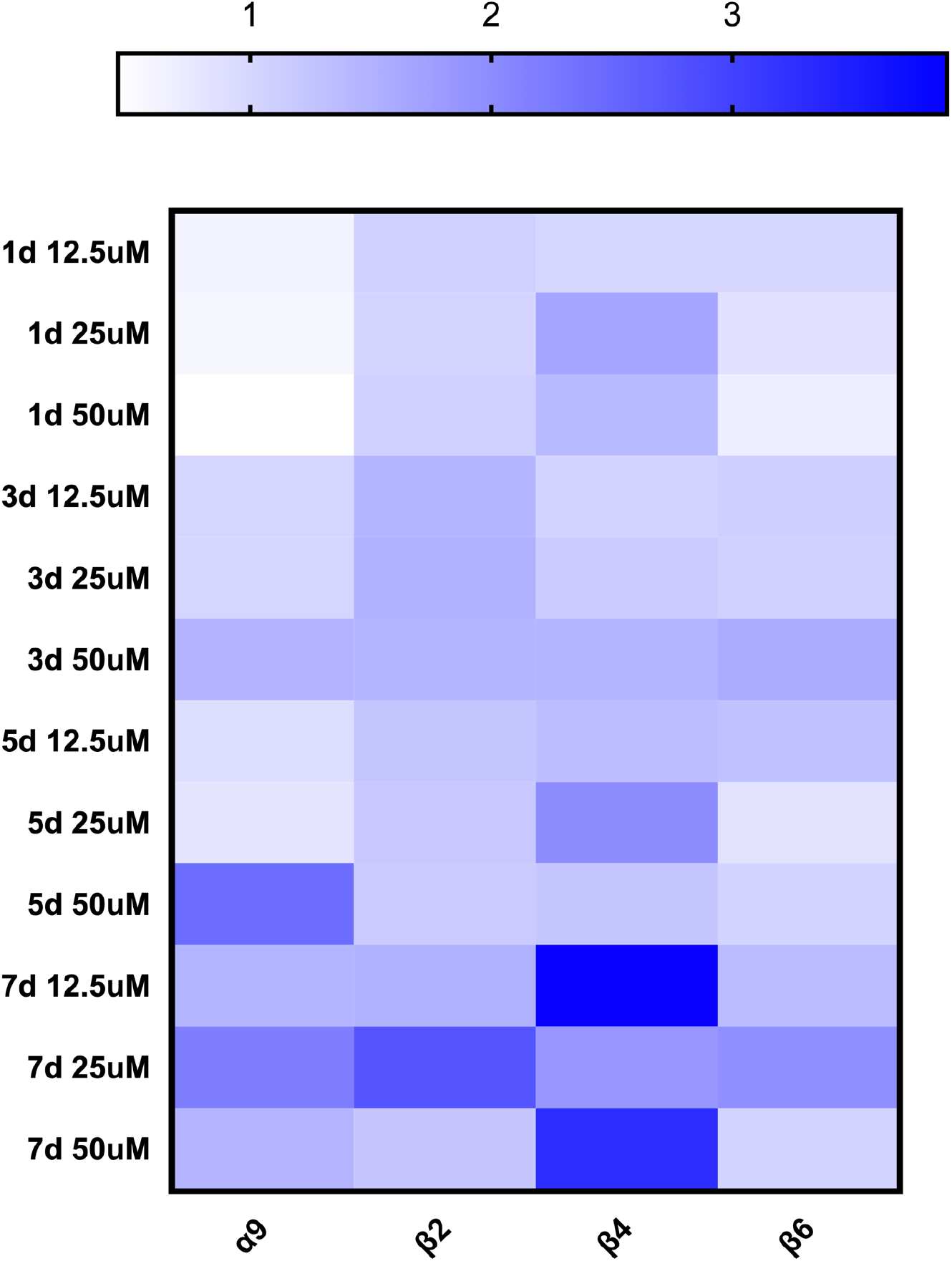
The expression level of tubulin genes mRNA under different administration time and concentration (level axis indicates gene name, vertical axis indicates relative fold change value).

As shown in Fig. 4, overall, the expression of β tubulin were decreased significantly than *α* tubulin, especially in day 1, day 3 and day 5. While, the *α* tubulin were changed slightly, in contrast to that of β tubulin.

**Fig 4.**
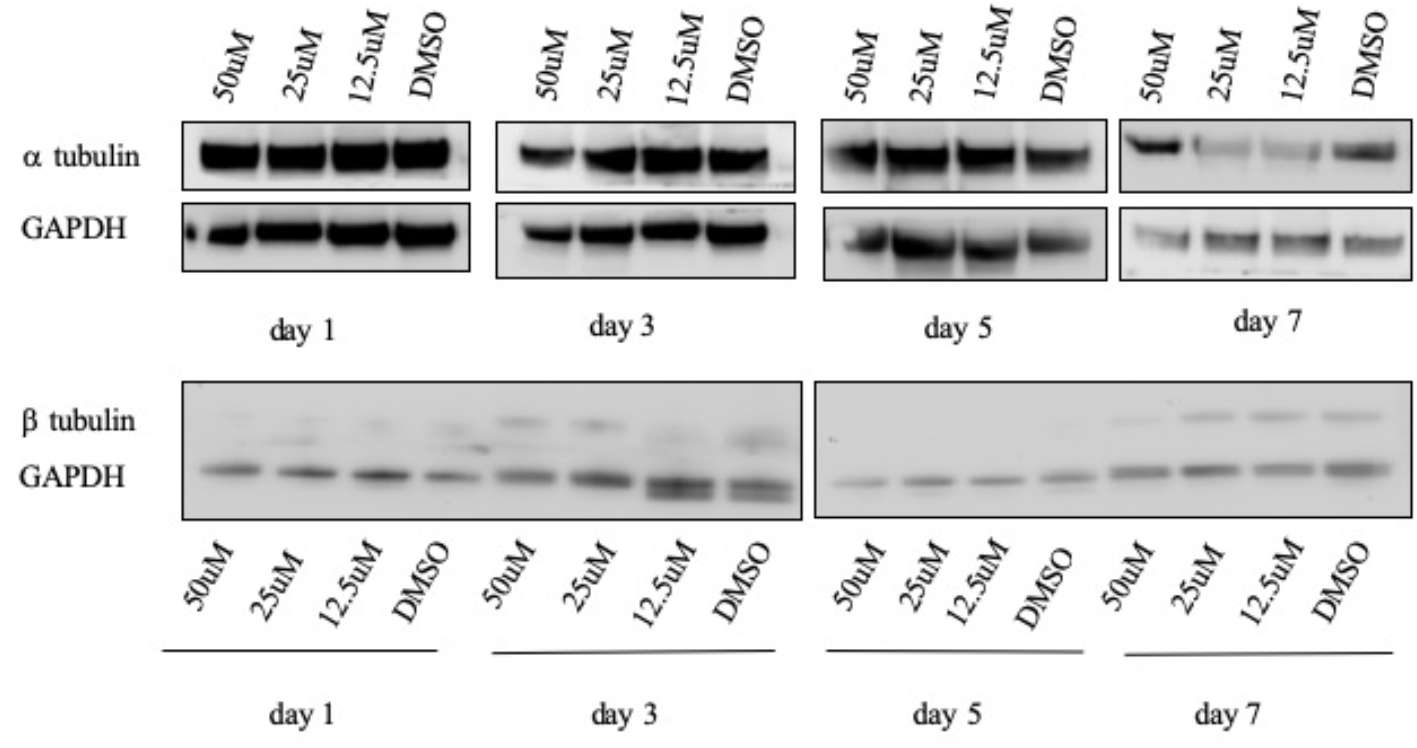
The protein expression of α tubulin and β tubulin under different administration time and concentration.

As shown in Fig. 5, in the untreated control samples, through TEM, we observed the intact well-defined flame cell, which has a special configuration of filled with approximately 60-80 cilia. Moreover, by careful observation of each cilium, we confirmed that these cilia were hexagonal in transverse section and comprised of microtubules, which arranged with nine doublets and two singlets, which we called the 9×2+2 structure, and each microtubule was hollow tubular structure (Fig. 5A, Fig. 6). A plasma membrane surrounds the cell body that extends through the tip of the tuft of cilia and is apparently in close interdigitation with the membrane of adjoining cells (33) (yellow arrows in Fig. 5 Aa).

**Fig 5.**
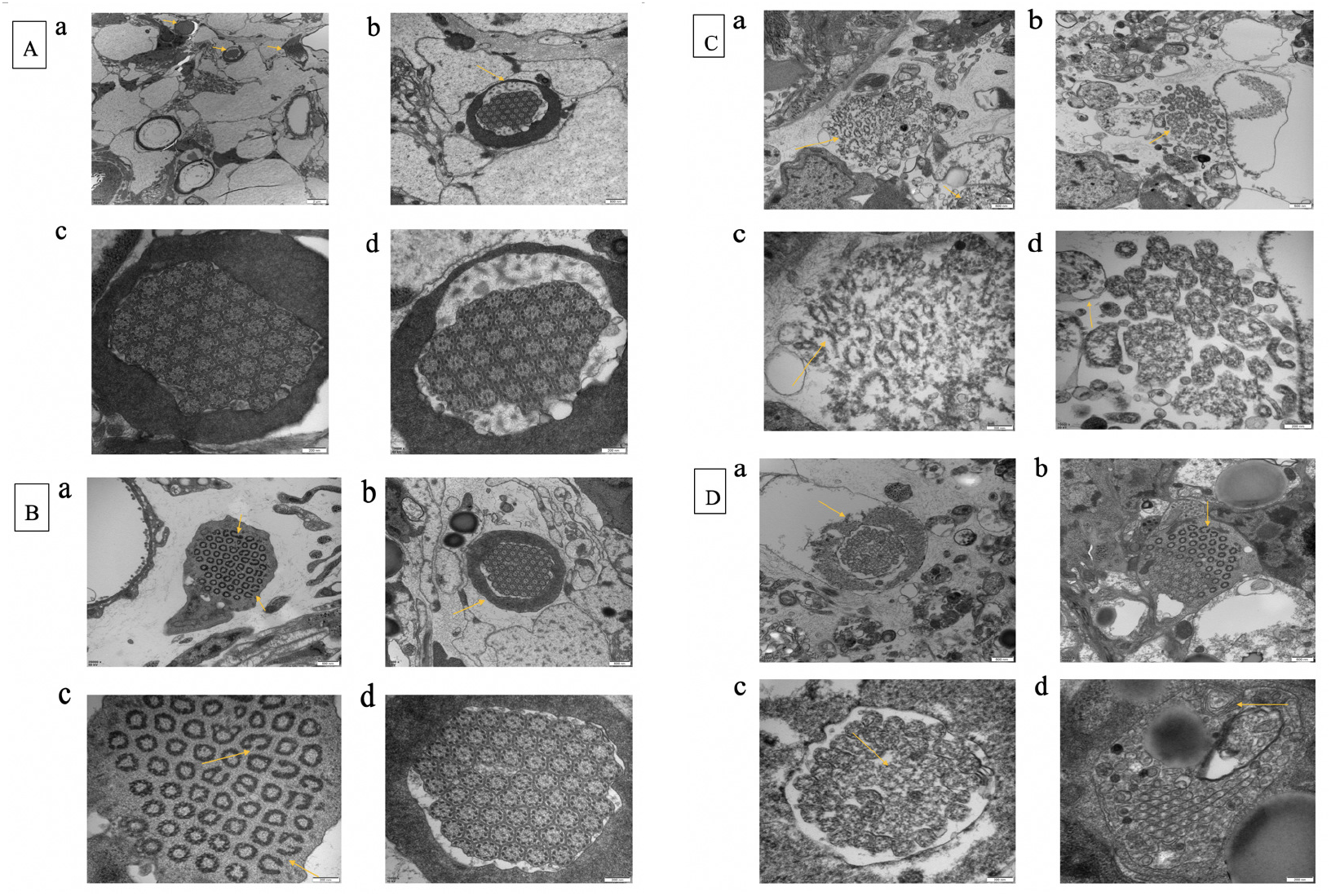
The ultrastructure changes of flame cell and microtubule in untreated control samples and MBZ 12.5uM, 25uM, 50uM treated groups through transmission electron microscope (correspond to A, B, C, D, respectively). Scale bars: Aa: 2um; Ab, Ba, Bb, Ca, Cb, Da, Db: 500 nm; Ac, Ad, Bc, Bd, Cc, Cd, Dc, Dd: 200nm. With concentration increased, the flame cell and microtubule were damaged increasingly apparent.

**Fig 6.**
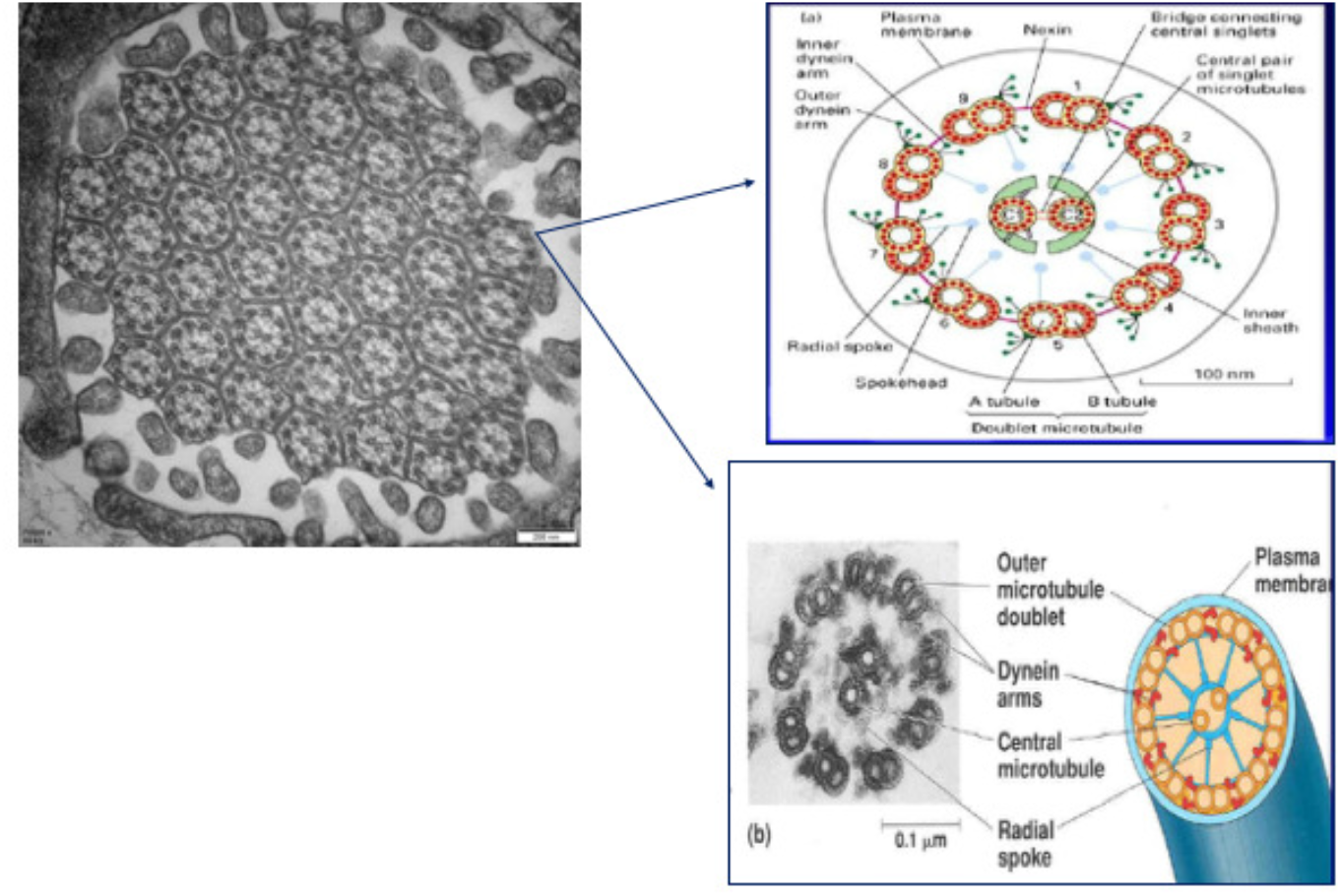
The ultrastructure of single cilium in the flame cell. (The right two photos quote from Wikipedia).

In Fig. 5A, 3 flame cells found together, and the microtubule structure of untreated control samples is complete (seen the yellow arrows in Fig. 5 Aa) and organized representative “9+2” structure regularly. More importantly, the flame cells membrane is integrated and around the compact dense tuft of cilia, constituted the circle cell body. By contrast, in the MBZ 12.5uM group, a few cilia are fractured, and a few microtubules are cracked (seen Fig. 5 Ba, Bc yellow arrow), but the whole structure of flame cell is visible, and the cilia are hexagonal in transverse section still on the whole (seen Fig. 5 Bb, Bd).

However, in MBZ 25uM group, the whole structure of flame cell is invisible, the flame cell membrane is ruptured, the fracture strength of cilia increased as well, the fracture cleft were extended, the arrange of tuft of cilia became scattered and twisted, the microtubule structure was even deformed (seen Fig. 5C yellow arrow). In addition, we observed there are small curled structure in Fig. 5 Ca. In MBZ 50uM group, the flame cell membrane is incomplete, few cilia are fractured, and few microtubules are cracked (seen fig. 5D yellow arrow). Furthermore, we observed some myelin sheath, presented alternately dark and bright lamellae concentric configuration (seen Fig. 5 Dd yellow arrow), the myelin sheath is a phenomenon which occurs when cell membranes are severely damaged. The myelin sheath phenomenon suggested that the cell membrane damage was much more serious than MBZ 25uM group and 12.5uM group.

### *In vitro* efficacy of RNAi against *E. multilocularis* metacestodes tubulin

As shown in Fig. 7A, the expression of tubulin genes *α 9, β*2, *β6* was reduced compared with the untreated sample, especially, the FC value of β2-2-treated sample is 9.01, suggested that the *β* 2 tubulin gene expressed reduced significantly. The difference of intergroup has statistical significance (p < 0.05). As shown in Fig. 7B, protein level detection of β tubulin and GAPDH was performed by western blotting, SDS-PAGE analysis showed a single purified β tubulin and GAPDH band, with a molecular weight of approximately 50 kDa, 35kDa. Western blotting analysis showed that the expression of β tubulin is down-regulated in siRNA-treated PSCs at day 15 post-electroporation. More generally, it suggests that the expression level of β tubulin appeared decreased with time, especially the β 2, β 6 tubulin.

**Fig 7.**
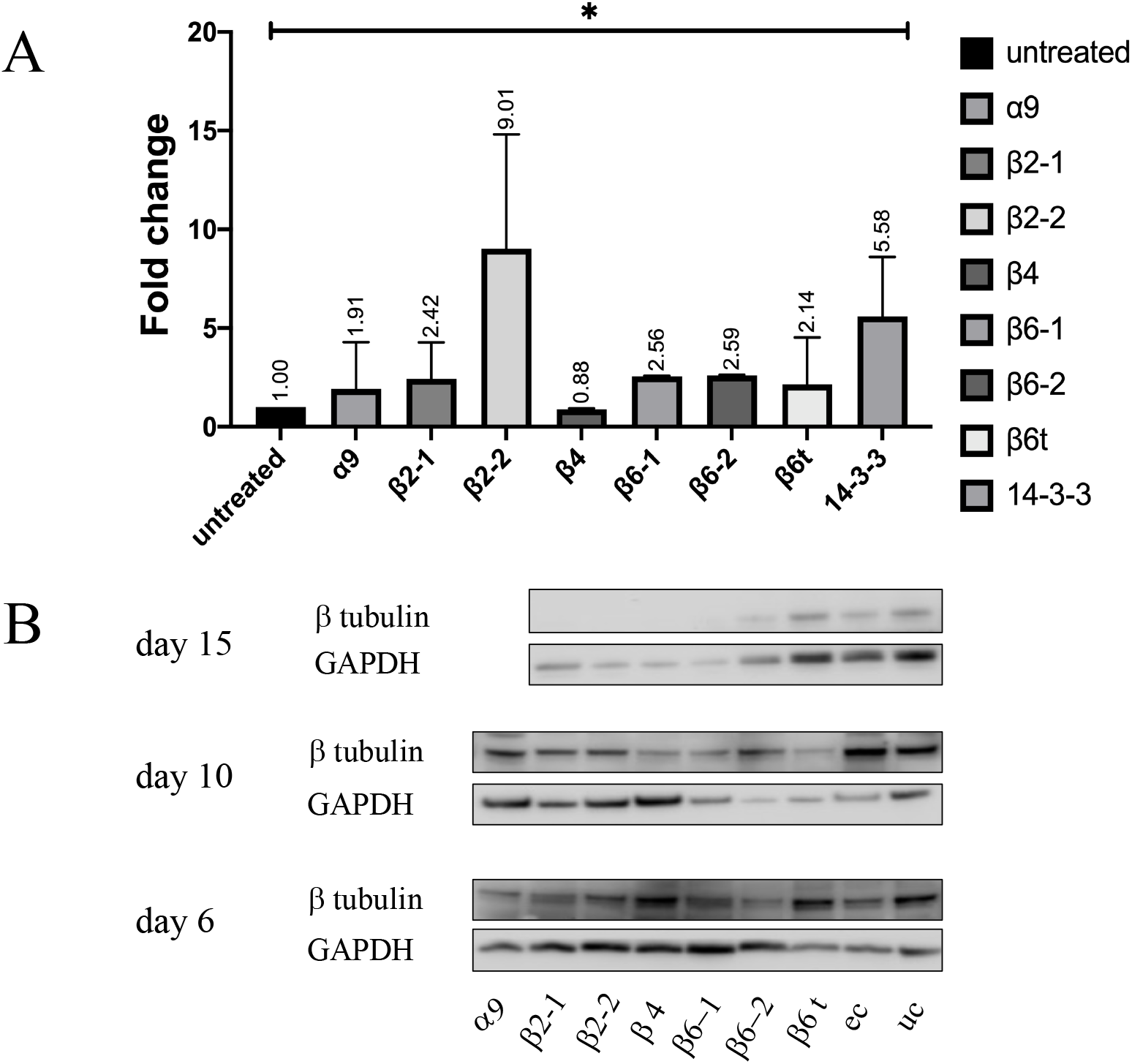
A showed RNAi effects of tubulin genes on mRNA levels postelectroporation 3 days in *E. multilocularis* PSCs. B showed RNAi effects of tubulin genes on protein levels post-electroporation 6, 10, 15 days in *E. multilocularis* PSCs. Bars represent the standard deviation. *p < 0.05.

In the untreated control samples, the flame cell is intact, it appears cupped (Fig. 8a), and we observed the complete flame cell and collective tubes complex (Fig. 8b, c, d), which is the excretory system of *E. multilocularis* PSCs. The bodies of flame cells are embedded in the parenchymal tissue, the cilia tufts extend to inside the protonephridial system tubules (collective tubes) (33). The protonephridial system is an excretory system that allows parasites to conserve water and eliminate salts. The cytoplasm of the flame cell extends to around, in order to interdigitation with other cells. It has abundant rough endoplasmic reticulum, and irregular vacuoles (actually is the cross section of collective tubes) in cytoplasm.

**Fig 8.**
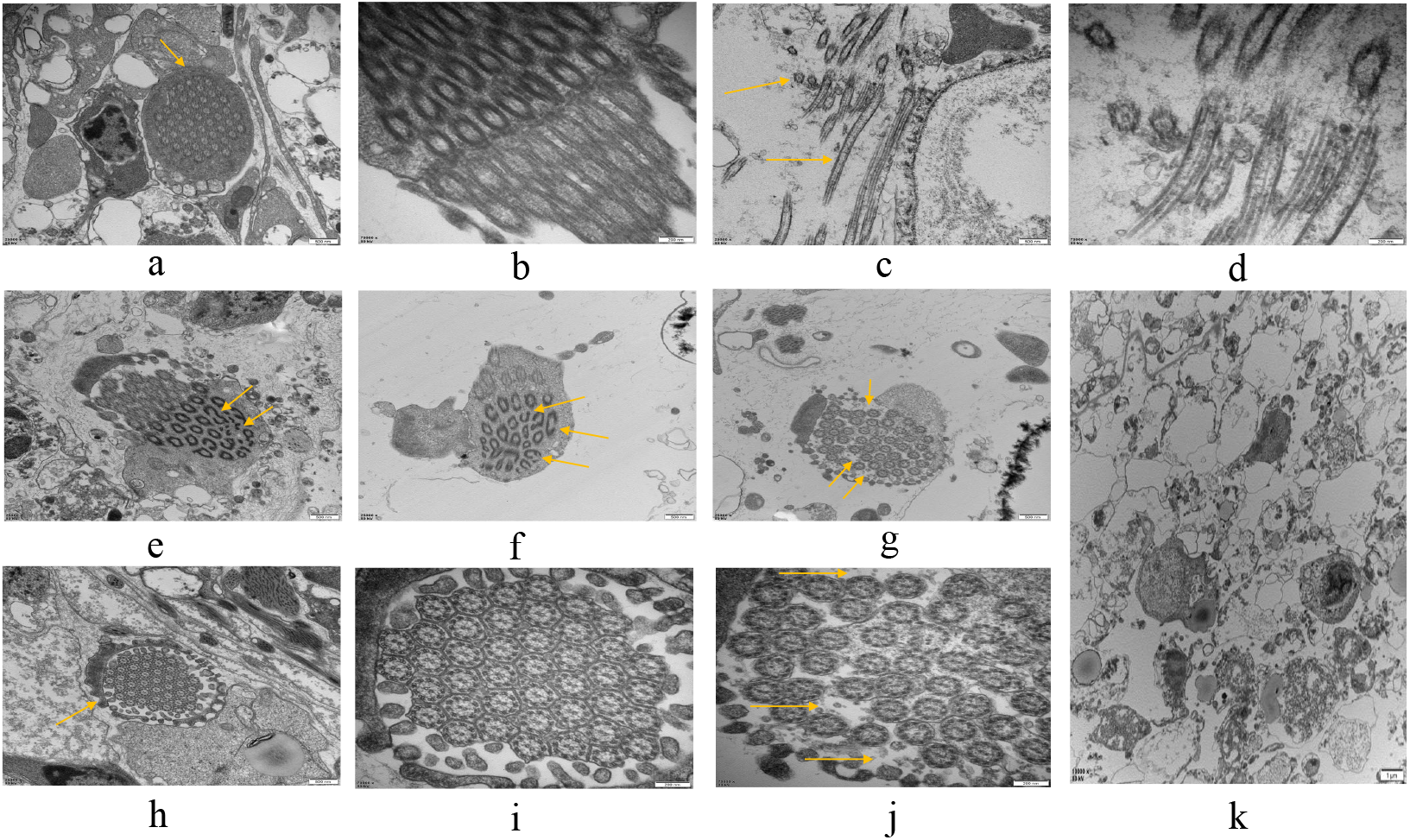
The ultrastructure changes of flame cell and microtubule in untreated control samples and *β 2-1, β 2*-2-treated samples through transmission electron microscope (TEM). Photo a, b, c, d reveals the untreated control samples, photo e-j reveals the *β 2*-1-treated samples, photo k reveals the *β 2*-2-treated samples. Scale bars: (b, d, i, j): 200nm, (a, c, e, f, g, h): 500 nm, k: 1um. Photo c, g, h corresponding to d, j, i, separately.

By contrast, in *β 2*-1-treated samples, a few cilia are fractured, some microtubules are cracked as well (yellow arrow in Fig. 8e, f), the microtubules are loose in cytoplasm, the whole structure of flame cell is visible, the cell membrane is complete (yellow arrow in Fig. 8g, j), and the cilia are hexagonal in transverse section still on the whole, many cytoplasm ecptoma (antler-like projects), nucleus at the bottom (seen Fig. 8h, i). In *β 2*-2-treated samples, we only observed some cell fragments, and vacuoles with different kinds and sizes (indistinction of collective duct) were observed, the nuclear membrane of tegument cell were swelled, the flame cell structure was not observed (seen Fig. 8k).

## DISCUSSION

Infection with the larval stage of the tapeworm *E. multilocularis* is a life-threatening disease in humans. Surgical resection of the involved liver segment and of metacestode lesions from other infected organs is indicated, but only one-third of patients with AE may benefit from curative liver resection, and the number is even lower in communities where AE is endemic and patients live in remote places and are late in seeking care (10, 34). Even after complete resection, recurrences have frequently been described (35). AE was therefore considered to be an incurable parasitic disease before the BZ derivatives MBZ and ABZ became available as chemotherapeutic agents (36). In a German study, with long-term MBZ chemotherapy with a mean duration of 3.9 years, the general clinical condition improved for 57% of the patients but remained unchanged for 22% of the patients. A survey of Alaskan patients (37) demonstrated that long-term chemotherapy with MBZ significantly increased the survival rate, from 25% for untreated patients to 90% for treated patients over a 10-year follow-up.

Unfortunately, despite the improvements in the chemotherapy of AE with BZ derivatives, complete erradication of the parasitic mass cannot be achieved in the majority of patients (16, 38), although one Alaskan study was more favorable, indicating that long-term application of MBZ may cause the death of the parasite (37). Our findings corroborate and strengthen previous results that have demonstrated concentration- and time-dependent effects of MBZ on PSCs tubulin, especially the β-tubulin.

Furthermore, to understand the β-tubulin action for *E. multilocularis* PSCs growths and development, we applied RNAi to knock down some high abundance tubulin genes to observe the development of PSCs, and demonstrated that knocked down *β 2* tubulin gene inhibited the development of flame cell, especially the development of ciliary tuft, thereby damaged the parasite structure, prevented parasites growth, and reduced the proliferation of *E. multilocularis* PSCs. Consequently, we concluded that the *β 2* tubulin gene maybe one of the crucial tubulin genes to form microtubule made up for the cilia tufts in flame cell.

RNA interference (RNAi) has been used successfully as a means of silencing specific gene expression to elucidate function and to assess the therapeutic value of candidate genes in helminths, including turbellarians (39), trematodes (40) and a monogenean (41), but there is relatively limited information on the use of the approach to determine gene function in cestodes. In this study, we used both soaking and electroporation to transfect siRNA into PSCs of *E. multilocularis* PSCs initially, and the soaking method was also useful for transfecting small RNA into PSCs. However, we found that fluorescence was only detected in the tegument of PSCs treated by soaking. By contrast, in parasites subjected to electroporation, strong fluorescence was detected in many calcareous corpuscles in PSCs after treatment, possibly because the electroporation shock may have temporarily opened channels on the tegumental surface of PSCs or destabilized membranes, resulting in the formation of membrane pores facilitating increased entry of siRNA or dsRNA into the worm tissues. This was same with the Hui Wang, et al study (28).

Flame cells are ciliated cells located within the basal matrix, at the neodermal tissue of cestoda. They are considered as the basic units of the protonephridial system of invertebrates. As for intestinal helminthes, to survive in the intestine or body cavities of a host, the parasite’s protonephridial system acts as an osmoconformer (33, 42). Parasites must also maintain osmotic pressure in their tissues within physiological limits against that of the host environment (43). The protonephridial system are the excretory systems with an important role that allow the parasites to conserve water and eliminate salts and survive in the intestine or body cavities of their hosts and because of that, they act like osmoconformers (43, 44). Flame cells mainly as key regulators of parasite metabolism, drug excretion and interaction with the host (45), and tubulin are the main cytoskeletal proteins that mediate proper function of flame cell cilia and maintain the excretory system (46). However, the flame cells were primarily affected at the level of the ciliary tuft, in association with the changes in tubulin, we found that altering tubulin expression and thus affecting the assembly and function of flame cells.

Our results have identified the MBZ action on four tubulin genes in molecular level, and how MBZ parasitostatic to *E. multilocularis* PSCs through impacting on the flame cell and microtubule, and decrease the *β 2*-tubulin gene expression changed ultrastructure and function of the flame cell, which facilitates exploitation of designing anthelmintic drugs that exclusively impair parasitic proteins which mediate cell signaling and pathogenic reproduction and establishment.

## ETHICS STATEMENT

The animal study was reviewed and approved by Animal care and all animal procedures were carried out in compliance with the Guidelines for the Care and Use of Laboratory Animals produced by the Shanghai Veterinary Research Institute. The study was approved by the Ethics Committee of the National Institute of Parasitic Diseases, Chinese Center for Disease Control and Prevention. The license number was IPD-2014-2.

## ACKNOWLEDGEMENTS

We gratefully acknowledge the technological help of Electron microscope Department, Shanghai Jiaotong University School of Medicine, during the study.

This study was supported by the General Program of Shanghai Municipal Health Commission Grant (No. 201940368).

We have no conflicts of interest to declare.

H.Z. and Y.T. conceived the ideas. Q.S. and L.H. collected the data. J.Z., B.J. and H.G. analyzed the data. Q.S. led the writing.

